# MicroRNA-21-3p regulation of NOX4 and VEGFA contributes to hemorrhage in cerebral cavernous malformations

**DOI:** 10.1101/2025.10.13.681886

**Authors:** Xin-Xing Guo, Zhong-Run Huang, Pei-Sheng Chen, Qi Li, Jia Li, Zhong-Song Shi

**Affiliations:** Department of Neurosurgery, Sun Yat-sen Memorial Hospital, Sun Yat-sen University, Guangzhou, China; RNA Biomedical Institute, Sun Yat-sen Memorial Hospital, Sun Yat-sen University, Guangzhou, China; Curtin Medical Research Institute, Curtin Medical School, Curtin University, Perth, Australia; Guangdong Province Key Laboratory of Brain Function and Disease, Sun Yat-sen University, Guangzhou, China

**Author notes:** Xin-Xing Guo, Zhong-Run Huang and Pei-Sheng Chen have contributed equally to this work. Correspondence: Zhong-Song Shi, Department of Neurosurgery, Sun Yat-sen Memorial Hospital, Sun Yat-sen University, 107 Yanjiang West Road, Guangzhou, Guangdong, China.

**Keywords:** Cerebral cavernous malformations, Endothelial cell, Pericyte, MicroRNA, Cerebral hemorrhage, Zebrafish

## Abstract

**Objective:** MicroRNAs regulate the brain vascular integrity and are involved in the lesion development of cerebral cavernous malformations (CCM). This study examines the role of microRNA-21-3p in CCM-related cerebral hemorrhage and its underlying mechanisms.

**Methods:** The expression of miRNA-21-3p and its target genes of NADPH oxidase 4 (NOX4) and vascular endothelial growth factor A (VEGFA) in brain microvascular endothelial cells (BMECs) and pericytes were assessed in cavernous malformation lesions of 20 sporadic CCM patients by fluorescence in situ hybridization. The association of their expression with hemorrhage manifestation was evaluated. Cell proliferation, permeability, reactive oxygen species (ROS), migration, tubule formation, and the expression of NOX4 and VEGFA were assessed in *CCM2* gene-depleted human BMECs and pericytes after miRNA-21-3p intervention. Cerebral hemorrhage, vascular permeability, vascular dilation, and angiogenesis after miRNA-21-3p intervention were evaluated in the *ccm2* gene-knockdown zebrafish.

**Results:** Decreased miRNA-21-3p and increased NOX4 and VEGFA were shown in BMECs and pericytes of the CCM lesions compared to peri-lesion normal vessels from epilepsy patients, which were also correlated with the presence of cerebral hemorrhage in CCM patients. Increasing miRNA-21-3p attenuated cell proliferation, permeability, ROS expression, cell migration, and tubule formation by targeting NOX4 and VEGFA in *CCM2* gene-depleted BMECs and pericytes. In vivo studies revealed that increasing miRNA-21-3p reduced cerebral hemorrhage, vascular permeability, vascular dilation, angiogenesis, and the overexpression of *nox4* and *vegfa* in *ccm2* gene-knockdown zebrafish.

**Conclusion:** MiRNA-21-3p can be a novel therapeutic target by regulating NOX4 and VEGFA, thereby stabilizing vascular integrity and reducing cerebral hemorrhage in CCM lesions.

## 1. Introduction

Cerebral cavernous malformations (CCMs) are brain vascular anomalies characterized by histological features of clusters of dilated, thin-walled blood vessels, which are clinically significant due to their potential to cause cerebral hemorrhage, significantly affecting the patient’s quality of life [1]. Surgical resection and stereotactic therapy remain challenges for lesions in eloquent areas and the brainstem; therefore, pharmacological therapy provides a potential venue for CCMs, given the emerging deeper understanding of these pathogeneses [2–6].

Familial CCMs predominantly arise from deletions and mutations in the *KRIT1* (*CCM1)*, *CCM2*, or *PDCD10* (*CCM3)* complex in endothelial cells by modulating key signaling pathways, including KLF2/4, endothelial-mesenchymal transition, RhoA/ROCK, and TLR4-like receptor signaling [7,8]. Somatic mutations of PIK3CA and MAP3K3 play a crucial role in sporadic CCMs. Functional loss mutations in the *CCM* complex and somatic mutations in PIK3CA enhance the mTOR signaling pathway in endothelial cells, forming a complex mechanism similar to cancer that promotes abnormal vascular growth and leads to the formation of CCMs. The mTORC1 inhibitor rapamycin can effectively inhibit the formation of cavernous malformations in mouse models [4,5,9].

Non-coding RNAs have a key role in the development of cerebral vascular malformations by regulating gene expression and influencing angiogenesis, inflammation, vascular integrity, apoptosis, and oxidative stress. MicroRNAs, as one of the important non-coding RNAs, have emerged as a potential value for diagnosis, prognostic prediction, and therapeutic use in CCMs [10–12]. Several microRNAs, including miRNA-181a-2-3p, miRNA-30c-2-3p, miRNA-3472a, miRNA-95-3p, miRNA-361-5p, miRNA-370-3p, and let-7b-5p, have been shown to link with CCMs [13–15]. A target site blocker that prevents the interaction of miRNA-27a with vascular endothelial cadherin can inhibit the increase in endothelial cell permeability and reduce lesion volume, as well as cerebral hemorrhage, in a mouse model of CCM [16].

More evidence suggests that pericytes and neuroglia are also involved in the CCM pathogenesis as non-endothelial cells. In CCM lesions, pericytes interacting with endothelial cells can activate vascular endothelial growth factor A (VEGFA)/VEGFR2 signaling, leading to increased angiogenesis and vascular permeability [5,17–20]. MicroRNAs, which can regulate gene expression and influence several CCM-associated pathological processes in both pericytes and endothelial cells, may serve as valuable targets for reducing hemorrhage in CCM disease.

MiRNA-21 has been identified as modulating apoptosis and inflammation involved in the response to oxidative stress and angiogenesis in ischemic stroke and glioblastoma [21,22]. Furthermore, miRNA-21 inhibits the excessive production of reactive oxygen species (ROS) induced by nicotinamide adenine dinucleotide phosphate oxidase 4 (NOX4), thereby suppressing the development of endothelial cell tumors [23]. In bovine ovarian granulosa cells, miRNA-21-3p regulates autophagy through downregulating VEGFA expression and inhibiting the PI3K/AKT/mTOR signaling pathway [24]. In CCM disease, the role of miRNA-21-3p in cerebral hemorrhage remains unclear. This study aims to investigate the relationship between miRNA-21-3p expression in brain microvascular endothelial cells (BMECs) and pericytes, and cerebral hemorrhage in CCM lesions. We further aim to clarify the mechanisms by which miRNA-21-3p regulates cerebral hemorrhage using both a human BMECs and pericytes model with *CCM2* gene depletion, as well as a zebrafish model with *CCM2* gene knockdown. By understanding the interactions between miRNA-21-3p and its target genes involved in oxidative stress and angiogenesis in both BMECs and pericytes, this study aims to identify a novel molecular mechanism that could be targeted to reduce cerebral hemorrhage, offering the potential of miRNA-21-3p as a therapeutic target in CCM disease.

## 2. Methods

### 2.1. Human CCM samples

The clinical, imaging, and histological data of 20 sporadic CCM patients in our hospital were analyzed in this study. These patients had received the initial surgical resection of the lesion without previous radiation therapy. They had the preoperative cranial MRI examination, excluding patients with extracranial CCM lesions, vascular malformation smaller than 1 cm, coexisting other neurological disorders, and mental illness.

The vascular malformation tissues were frozen in liquid nitrogen, immediately resectioned, and stored at −80 °C. Then, RNA was extracted to detect the expression of miRNA-21-3p, *CCM1*, *CCM2*, and *CCM3* in the cavernous malformation lesions using RT-PCR. The CCM tissue specimens were embedded in paraffin and sectioned for fluorescence in situ hybridization staining to detect the expression of miRNA-21-3p, NOX4, and VEGFA in the endothelial cells and pericytes of vascular malformation lesions. The lesions from five patients with temporal lobe epilepsy (TLE) and glioblastoma multiforme (GBM) who received their first surgical resection without radiation therapy history were used as the control group. The study was reviewed and approved by the Ethics Committee in our institution.

### 2.2. Cerebral hemorrhage on MR imaging

CCM patients underwent preoperative cranial MR examination on a 3.0T magnetic resonance scanner (Achieva TX, Philips Healthcare) with scanning sequence including T1-weighted imaging, T2-weighted imaging, T2 fluid-attenuated inversion recovery, 3D time-of-flight angiography, and susceptibility-weighted imaging. The number of CCM lesions, location, size, and concomitant cerebral hemorrhage were analyzed. The hemorrhage manifestations of CCMs on MR have different signals around the vascular malformation lesion according to the acute, subacute, and chronic bleeding period. In this study, the hemorrhage manifestations of CCMs were defined as acute or subacute bleeding signals with a thickness greater than 4mm around the CCM lesion on T2-weighted imaging, as in the previous study [25].

### 2.3. Fluorescent in situ hybridization and immunofluorescent

To determine the cell-type specific expression of miRNA-21-3p, NOX4, and VEGFA, we combined fluorescent in situ hybridization (FISH) with immunofluorescent, as previously described [26,27]. In brief, paraffin-embedded CCM sections were prehybridized for 2 hours at 50°C, followed by hybridization with a digoxin-labeled miRNA-21-3p probe (QIAGEN, Germany) overnight at the same temperature. The sections were then washed three times with SSC buffer at 37°C. Next, biotinylated mouse anti-digoxin antibody was applied and incubated for 2 hours at room temperature, followed by rinsing with PBS. Subsequently, Streptavidin-Biotin System Fluorescein Isothiocyanate was added and incubated for 30 minutes at 37°C. To specifically label endothelial cells and pericytes, rabbit anti-CD31 antibody (1:50, ab215912, Abcam) and rabbit anti-PDGFR-β antibody (1:50, ab51745, R&D Systems) were added, followed by rabbit anti-NOX4 antibody (1:50, ab133303, Abcam) and rabbit anti-VEGFA antibody (1:50, ab51745, Abcam), incubating overnight at 4°C. After rewarming to room temperature for 45 minutes, the sections were washed with PBS for 5 minutes. Finally, a fluorescent secondary antibody was added dropwise in the absence of light and incubated for 1 hour at 37°C. Subsequently, 4’,6-diamidino-2-phenylindole (DAPI) was applied to stain the nuclei for 10 minutes. Images were captured using a Zeiss LSM800 confocal microscope, with the expression levels of miR-21-3p, NOX4, and VEGFA quantified as average fluorescence intensity using ImageJ.

### 2.4. Human brain microvascular endothelial cells and pericytes

Primary human BMECs and brain pericytes were provided by Cell System (ACBRI 376, ACBRI 498), cultured in Complete Classic Medium (Cell systems,4Z0-500) or pericyte medium with 2% fetal bovine serum and supplements (Shanghai QiDa, P4001), and used at passage 3.

### 2.5. siRNA knockdown and miRNA-21-3p intervention in *CCM2* gene-depleted human BMECs and pericytes

Small interfering RNA (siRNA) was used to knock down the *CCM2* gene in the human BMECs and pericytes with Lipofectamine 3000 Transfection Kit (Thermo Fisher Scientific, L3000-015) as previous studies [16]. The cells were maintained overnight, and the culture medium was changed the next day. For transfection with miRNA-21-3p overexpression or knockdown and *NOX4* or *VEGFA* knockdown in the *CCM2* gene-depleted cells, the siRNA transfection medium was maintained for 6-8 hours. Then, cells were transfected with miRNA-21-3p mimic, miRNA-21-3p inhibitor, microRNA negative control, siRNA *NOX4* with Lipofectamine 3000, or siRNA *VEGFA* with Lipofectamine 3000. Cells were maintained overnight and changed to culture medium the next day. Human BMECs and pericytes were collected 24 or 48 hours after transfection for further RNA assay, protein assay, cell proliferation assay, cell migration assay, tube formation assay, permeability assay, and ROS measurement. The optimal concentrations of the miRNA-21-3p mimics and miRNA-21-3p inhibitors were determined as 50nM and 100nM, respectively. The optimal siRNA sequences for *CCM2*, *NOX4*, and *VEGFA* were determined and shown in Supplementary Table S1. The miRNA-21-3p mimics and inhibitors were synthesized by RiboBio (Guangzhou, China). siRNA genes were synthesized by Gene Pharma (Suzhou, China).

### 2.6. Zebrafish model with *ccm*2 gene knockdown

Both AB wild-type and Tg (Fli1:eGFP; Gata1:DsRed) transgenic zebrafish were utilized in this study. All the zebrafish were bred and housed in a pathogen-free environment in a vivarium approved by the Laboratory Animal Care at our University. All animal experiments and protocols were approved by the Ethics Committee in our institution and conducted according to relevant guidelines and regulations. Knockdown and overexpression experiments were performed by injecting morpholino oligonucleotides (MO) or mimics into one-cell stage embryos.

The MOs of *ccm*2 and miRNA-21-3p were synthesized by Gene Tools (Philomath, OR, USA). The miRNA-21-3p mimics were synthesized by Gene Pharma (Suzhou, China). The amounts used for microinjection were 200 µM *ccm2* MO, 200 µM control MO, 100 µM miRNA-21-3p MO, 10 µM miRNA-21-3p mimic, and 10 µM miRNA negative control. In the rescue experiment, *ccm*2 MO was injected simultaneously with miRNA-21-3p mimic. The sequence of *ccm*2 MO, miRNA-21-3p MO, and miRNA-21-3p mimics used for zebrafish are shown in Supplementary Table S2.

### 2.7. In vivo confocal imaging for zebrafish model

To block the pigment formation of zebrafish at 1 day post-fertilization (dpf), 0.003% 1-pheny1-2-thiourea (P7629, Sigma) was added to 10% Hanks’ solution. Then, 2 mg/ml pronase (3115844001, Roche) was added to 10% Hanks’ solution to remove the zebrafish chorion to unify the cerebrovascular development time of zebrafish. The zebrafish embryos at 3-dpf were anesthetized by incubating in 10% Hanks’ solution containing a final concentration of 0.003% tricaine (E10521, Sigma) and embedded in 1% low-melting agarose (16520050, Invitrogen), then were subjected to in vivo imaging studies with an Olympus laser confocal microscope.

### 2.8. Detection of cerebral hemorrhage, brain vascular integrity, cerebrovascular dilation, and angiogenesis in zebrafish model

The cerebral hemorrhage, brain vascular integrity, cerebrovascular dilation, and angiogenesis in 3 dpf transgenic zebrafish embryos with *ccm*2 gene knockdown after miRNA-21-3p intervention were determined by in vivo confocal imaging. The fluorescence images were analyzed with ImageJ. The blood cell leakage into the brain of each zebrafish larva was marked, and the cerebral hemorrhage area was calculated using ImageJ [28,29]. The area of the primordial midbrain channel (PMBC) and the vessel branch points of the subintestinal vein (SIV) were quantified to determine the changes in cerebrovascular dilation and angiogenesis in zebrafish [30,31].

A 10-nL DAPI with a concentration of 0.8 mg/mL (C1002, Beyotime) was microinjected into the common cardinal vein of zebrafish at 3dpf. At 30 minutes after microinjection, fluorescence images were taken by a confocal microscope. The number of DAPI-positive parenchymal nuclei located outside the cerebral vessel was quantified to determine the brain vascular integrity in zebrafish using ImageJ [28,32].

### 2.9. Quantitative Real-Time PCR

The total RNA was extracted from human CCM lesions, BMECs, pericytes, and zebrafish tissue using Trizol (Thermo Fisher Scientific). The human brain total RNA was provided by Takara Bio (636530). PrimeScript Reverse Transcription Kit (RR036A for mRNA, RR037A for microRNA, Takara Bio) was used for cDNA synthesis. Quantitative Real-time PCR was performed on the Applied Biosystems PCR System with TB Green Premix Ex Taq II (RR820A, Takara Bio). U6 and β-actin were housekeeping controls for microRNA and mRNA expression analysis. The microRNA and mRNA expression were calculated using the 2-ΔΔCT method. The primer sequences of miRNA-21-3p and mRNA (*CCM1*, *CCM2*, *CCM3*, *NOX4*, and *VEGFA*) are synthesized by RiboBio and shown in Supplementary Table S3.

### 2.10. Western blot analysis

Proteins were isolated and quantified following standard protocols. A Western blot assay was conducted in accordance with established procedures. Densitometric analysis was performed using ImageJ software. The following antibodies were employed for Western blotting: goat anti-CCM2 (1:1000, SAB2500214, Sigma), rabbit anti-NOX4 (1:1000, ab133303, Abcam), rabbit anti-VEGFA (1:1000, ab51745, Abcam), and rabbit anti-β-actin (1:1000, 12620, Cell Signaling Technology). HRP-linked secondary antibodies included goat anti-rabbit IgG HRP (1:1000, 7074, Cell Signaling Technology) and rabbit anti-goat IgG HRP (1:1000, BL004A, Biosharp). The expression of β-actin served as an internal reference.

### 2.11. Cell proliferation assay

The CCK8 kit (CK04, DOJINDO, Japan) was employed to assess cell proliferation. Human BMECs and brain pericytes (1×10^3^ cells/well) were seeded into 96-well plates and incubated overnight at 37°C. Following transfection, 10 µl of the CCK8 solution was added to each well, and the mixture was incubated for 1 hour. Subsequently, the absorbance at 450 nm was measured using a microplate reader (Tecan, Switzerland).

### 2.12. Cell migration assay

Cell migration assay was conducted using Transwell chambers (353096, Corning, USA). Following transfection, 2×10^4^ cells were seeded in the upper chamber, and a medium without fetal bovine serum (FBS) was added. In the lower chamber, a medium containing 20% FBS was placed. After a 12-hour incubation period, the chambers were fixed with methanol and stained with 0.1% crystal violet for 15 minutes. The cells that migrated through the membrane were observed using a microscope (Olympus, Japan) and analyzed with ImageJ.

### 2.13. Tube formation assay

After transfection, human BMECs were inoculated at a density of 10,000 cells per well in 24-well plates pre-coated with Matrigel matrix (356234, Corning, USA) and cultured at 37°C for 12 hours. The formed tubules were monitored using a microscope (Olympus, Japan), and the number of tubules and branch lengths were quantified with the Angiogenesis Analyzer for ImageJ.

### 2.14. Transwell permeability assay in endothelial cells

The permeability of cell monolayers to sodium fluorescein (Sigma, USA) was measured as described previously [16]. Briefly, pre-transfected cells were plated and allowed to achieve confluence on the upper of the Transwell membrane.

Subsequently, 200 µl of sodium fluorescein (46960, Sigma, USA) was added to the upper chamber at 100 µg/ml and incubated for 30 minutes. The fluorescence of the lower chamber medium was then measured using an enzyme labeling instrument (Tecan, Switzerland) with an excitation wavelength of 460 nm and an emission wavelength of 515 nm.

### 2.15. Reactive oxygen species measurement

DCFH-DA (D6883, Sigma, USA) and DHE (D7008, Sigma, USA) were utilized to detect hydrogen peroxide and superoxide anions, respectively, as previously described [33]. Human BMECs and pericytes were seeded at a density of 1×10^5^ cells per well in 6-well plates and cultured overnight for transfection. Subsequently, the cells were incubated with 10 µM of DCFH-DA or DHE for 30 minutes at 37°C. After repeated rinsing several times, the cells were placed under a fluorescence microscope (Olympus, Japan) for observation, and the average fluorescence intensity was subsequently analyzed with ImageJ.

### 2.16. Luciferase reporter assay

A dual luciferase assay was performed to identify the target genes of miRNA-21-3p. HEK 293T cells were transfected with control and miRNA mimics for 48 hours. Following this, cells were transfected with the pGL3-NOX4-3’-UTR firefly luciferase expression construct (Genechem, Shanghai, China) or a construct carrying a mutation of the predicted miRNA-21-3p binding site in pGL3-NOX4-3’-UTR, together with the Renilla luciferase pRL-CMV expression construct using Lipofectamine 3000. Cell lysates were assayed using a dual luciferase reporter assay kit (Promega, USA). The data are presented as the ratio of firefly to Renilla luciferase activity.

### 2.17. Statistical analysis

Data are presented as mean ± SEM and were analyzed using SPSS 31.0 software. One-way ANOVA or independent-samples t-tests were employed to examine normally distributed data. The Mann-Whitney non-parametric test was utilized for non-normally distributed data. *P* < 0.05 was considered statistically significant.

## 3. Results

### 3.1. Decreased miRNA-21-3p expression and increased expression of its target gene of NOX4 and VEGFA in human CCMs

Twenty sporadic CCM patients included 9 males and 11 females, aged 2 to 72 years. Initial symptoms included epilepsy in 7 patients (35%), hemorrhage-related symptoms in 5 patients (25%), focal neurological deficits in 2 patients (10%), non-specific symptoms in 5 patients (25%), and asymptomatic physical examination in 1 patient (5%). The size of CCM lesions ranged from 1.1 to 4.0 cm, with 16 supratentorial lesions and 4 in the brainstem or cerebellar hemisphere. The clinical and cranial MR imaging data are presented in Table 1.

**Table 1.**
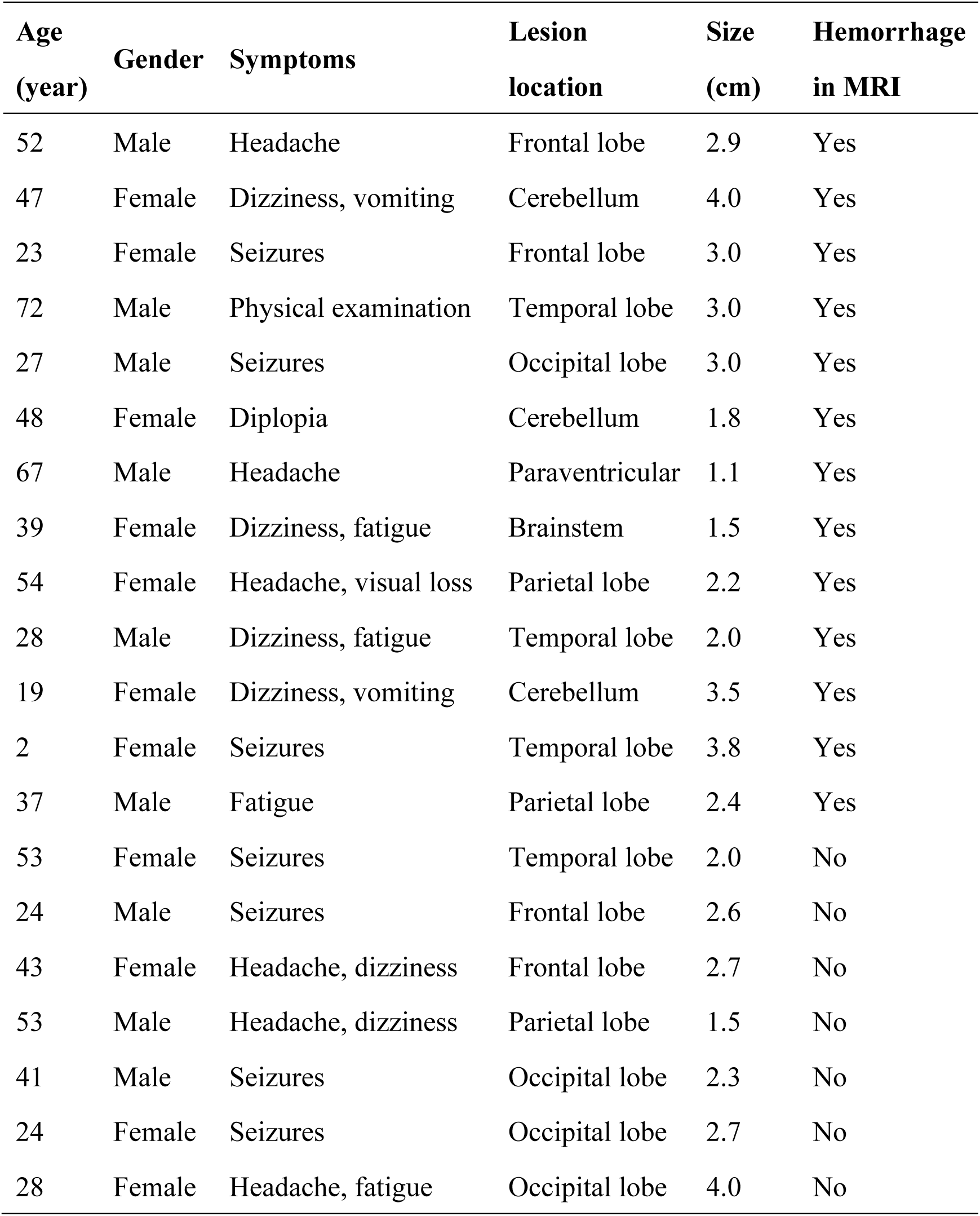
The clinical and cranial MR imaging data in 20 sporadic cerebral cavernous malformation patients.

MiRNA-21-3p expression was consistently positive in endothelial cells (ECs) of pseudonormal TLE and abnormal GBM vessels, and its expression in CCM lesion ECs was discontinuous with slight positive staining. MiRNA-21-3p expression in ECs of CCM lesion was significantly reduced compared to pseudonormal TLE and abnormal GBM vessels (Figures 1A and 1B). NOX4 and VEGFA expression in ECs of pseudonormal TLE vessels and abnormal GBM vessels showed slight positive staining, and its expression in ECs of CCM lesion was significantly positive. We found a significantly increased NOX4 and VEGFA expression in ECs of CCM lesion compared to TLE and GBM vessels (Figures 1A and 1B).

**Figure 1.**
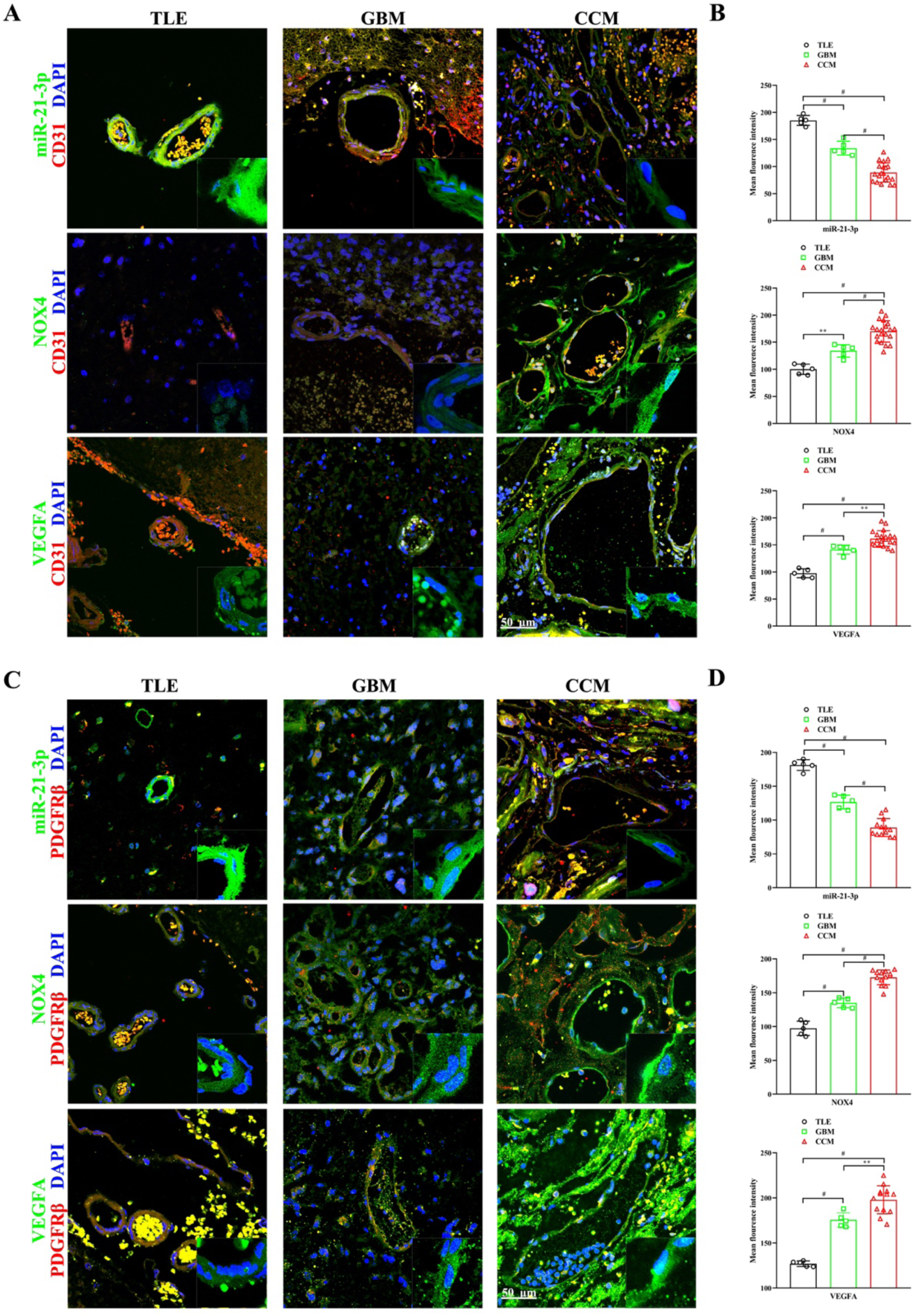
The expression of miR-21-3p was decreased in endothelial cells and pericytes of human cerebral cavernous malformations, while the expressions of NOX4 and VEGFA were elevated. A and B, The expression of miR-21-3p (green) in endothelial cells (red) of temporal lobe epilepsy (n=5), glioblastoma multiforme (n=5) and CCM (n=20) decreased gradually, but the expression of NOX4 and VEGFA (green) gradually increased. C and D, The expression of miR-21-3p (green) in pericytes (red) of temporal lobe epilepsy (n=5), glioblastoma multiforme (n=5) and CCM (n=13) decreased gradually, but the expression of NOX4 and VEGFA (green) gradually increased. Single staining for miR-21-3p, NOX4 and VEGFA of boxed areas is shown in the lower right. Nuclei are blue by DAPI. **P<0.01; ^#^P<0.001.

In pericytes, miRNA-21-3p, NOX4, and VEGFA expression followed a similar pattern. We found a significant decrease in miRNA-21-3p and an increase in NOX4 and VEGFA in pericytes of CCM lesion, mirroring the pattern in ECs (Figures 1C and 1D).

MiRNA-21-3p and *CCM1* and *CCM2* mRNA levels were significantly reduced in CCM lesions compared to normal brain tissue. There was no significant change in *CCM3* mRNA expression (Supplementary Figure S1).

### 3.2. Decreased miRNA-21-3p expression and increased NOX4 and VEGFA expression with cerebral hemorrhage in human CCMs

Thirteen of 20 CCM patients showed manifestations of cerebral hemorrhage on MRI (Figure 2A). In these patients, miRNA-21-3p expression in ECs was significantly reduced, while NOX4 and VEGFA expression levels were significantly increased (Figures 2B and 2C). Similar patterns were observed in pericytes (Figures 2D and 2E). There was no correlation between miRNA-21-3p, NOX4, and VEGFA expression levels and epilepsy manifestations in CCM patients.

**Figure 2.**
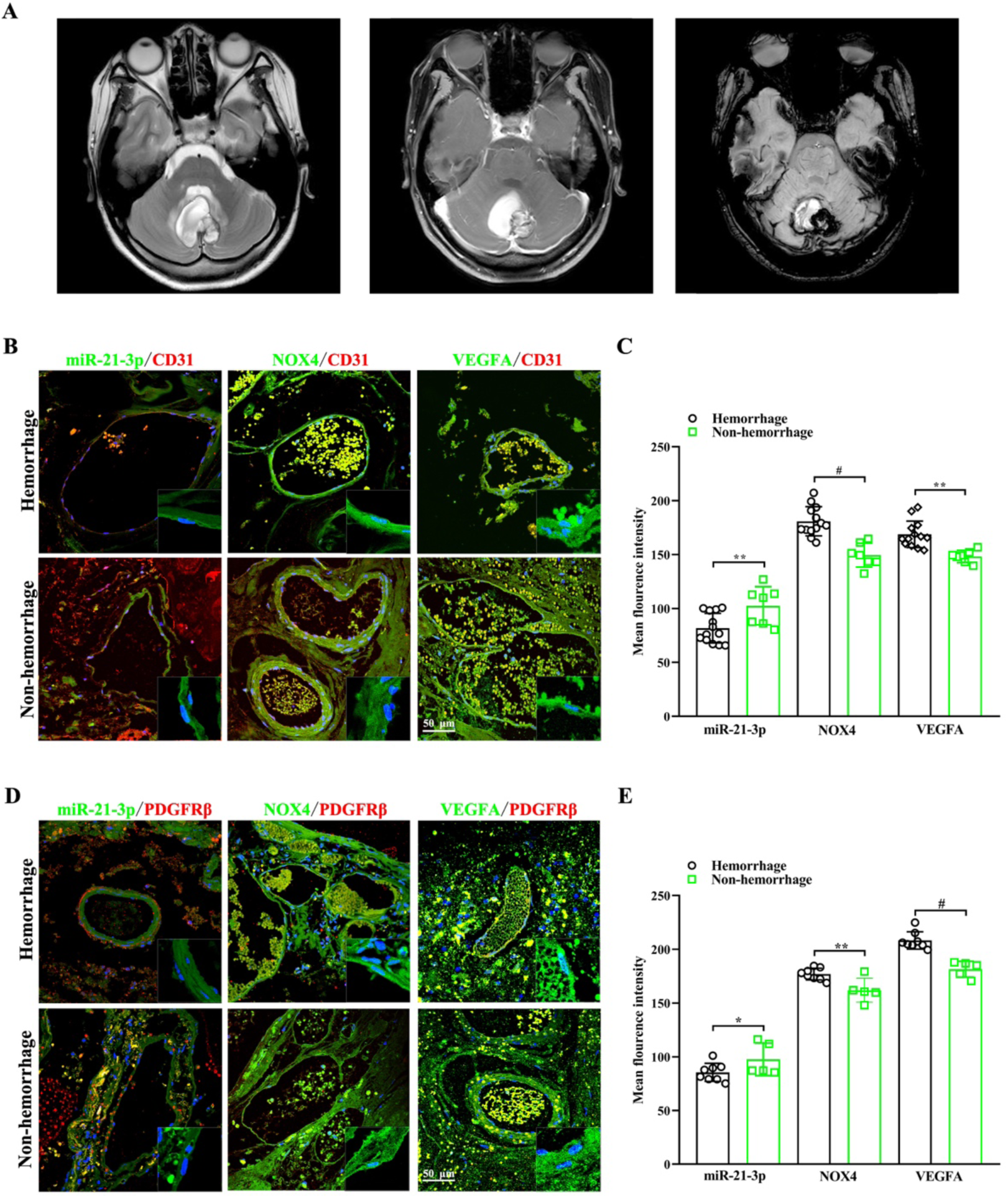
In the cerebral hemorrhage group of cerebral cavernous malformations, the expression of miR-21-3p in endothelial cells and pericytes was downregulated, while the expressions of NOX4 and VEGFA were upregulated compared to the non-hemorrhagic group. A, Typical cerebral hemorrhage images of cranial MRI of CCM patients, including T2-weighted imaging sequence (left), T1-weighted imaging enhancement sequence (middle), and susceptibility-weighted imaging sequence (right). B and C, Compared with the non-hemorrhage group (n=7), the expression of miR-21-3p (green) was decreased in the endothelial cells (red) of CCM lesions with cerebral hemorrhage (n=13) on MRI, but the expression of NOX4 (green) and VEGFA (green) increased. D and E, Compared with the non-hemorrhage group (n=5), the expression of miR-21-3p (green) was decreased in the pericytes (red) of CCM lesions with cerebral hemorrhage (n=8) on MRI, but the expression of NOX4 (green) and VEGFA (green) increased. Single staining for miR-21-3p, NOX4 and VEGFA of boxed areas is shown in the lower right. Nuclei are blue by DAPI. *P<0.05; **P<0.01; ^#^P<0.001.

### 3.3. MiRNA-21-3p regulated NOX4 and VEGFA expression in a human BMEC model with *CCM2* gene depletion

Given the decreased *CCM2* expression in human CCM lesions, we used siRNA to knockdown *CCM2* gene in a human BMEC model for an in vitro study. MiRNA-21-3p expression was significantly reduced after *CCM2* knockdown and could be modulated with miRNA-21-3p mimics or inhibitor (Figure 3A).

**Figure 3.**
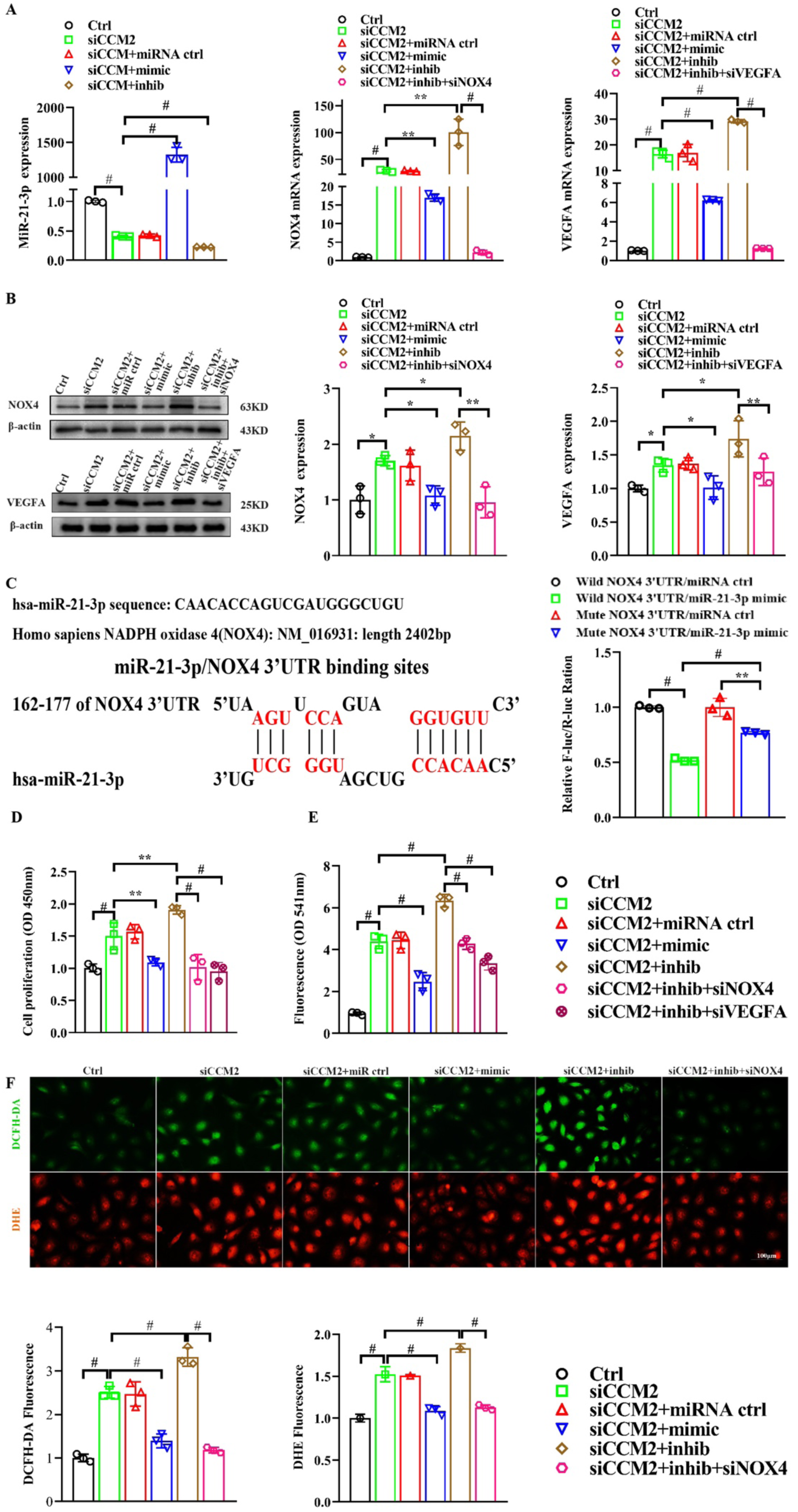
MiR-21-3p inhibits cell proliferation, endothelial permeability, and reactive oxygen species generation by suppressing the expression of the target genes NOX4 and VEGFA in *CCM2*-deficient human brain microvascular endothelial cells (BMECs). A, RT-PCR analysis of miR-21-3p, NOX4, and VEGFA mRNA expression in *CCM2*-deficient human BMECs transfected with miR-21-3p mimic and inhibitor. B, Western blot analysis of NOX4 and VEGFA protein in *CCM2*-deficient human BMECs transfected with miR-21-3p mimic and inhibitor. C, The putative binding sites of human miR-21-3p and *NOX4* 3’UTR were obtained from the Targetscan database. The dual luciferase assay indicated that *NOX4* was a direct target gene of miR-21-3p. D, *CCM2*-deficient human BMECs transfected with miR-21-3p mimic and inhibitor were subjected to the CCK-8 assay after transfection. E, Effect of miR-21-3p mimic and inhibitor on the permeability of the *CCM2*-deficient human BMECs monolayers to sodium fluorescein by in vitro permeability assay. F, Detection of superoxide by DHE and hydrogen peroxide by DCFH-DA in *CCM2*-deficient human BMECs transfected with miR-21-3p mimic and inhibitor. Mean ± SD are provided (n=3). *P<0.05; **P<0.01; ^#^P<0.001.

NOX4 and VEGFA expression levels were significantly increased after *CCM2* knockdown. MiRNA-21-3p mimic intervention significantly inhibited NOX4 and VEGFA upregulation, while miRNA-21-3p inhibitor intensified it. Combined miRNA-21-3p inhibitor and NOX4 or VEGFA siRNA interventions weakened the upregulation effects, indicating that miRNA-21-3p regulates NOX4 and VEGFA in *CCM2* gene-depleted endothelial cells (Figures 3A and 3B).

### 3.4. NOX4 and VEGFA as direct target genes for miRNA-21-3p

Bioinformatics tools predicted that *NOX4* is a target gene for miRNA-21-3p. Dual luciferase reporter assays confirmed that miRNA-21-3p binds to sites on *NOX4* 3’UTR, inhibiting its expression (Figure 3C). Previous study and dual luciferase assay also demonstrated that miRNA-21-3p directly targets *VEGFA* by binding to specific sites on its 3’UTR [24].

### 3.5. MiRNA-21-3p inhibits cell proliferation, permeability, and ROS by regulating NOX4 and VEGFA in a human BMEC model with *CCM2* gene depletion

Knocking down *CCM2* gene in BMEC led to increased cell proliferation, which miRNA-21-3p mimic intervention inhibited, while miRNA-21-3p inhibitor had the opposite effect. Combined miRNA-21-3p inhibitor with *NOX4* or *VEGFA* siRNA weakened the proliferation effects (Figure 3D). BMEC permeability increased following *CCM2* knockdown. MiRNA-21-3p mimic intervention restrained this rise, while miRNA-21-3p inhibitor intensified it. Combined miRNA-21-3p inhibitor with *NOX4* or *VEGFA* siRNA weakened BMEC permeability effects (Figure 3E). These results suggested that miRNA-21-3p attenuated proliferation and permeability in *CCM2*-depleted BMEC by regulating NOX4 and VEGFA.

Hydrogen peroxide and superoxide anion levels significantly increased after *CCM2* knockdown. MiRNA-21-3p mimic intervention inhibited this rise, while miRNA-21-3p inhibitor intensified it. Combined miRNA-21-3p inhibitor with *NOX4* siRNA weakened the upregulation effects, suggesting miRNA-21-3p attenuate ROS expression by regulating NOX4 (Figures 3F).

### 3.6. MiRNA-21-3p reduces angiogenesis by regulating VEGFA in a human BMEC model with *CCM2* gene depletion

Endothelial cell migration increased after *CCM2* knockdown. MiRNA-21-3p mimic significantly inhibited migration, while miRNA-21-3p inhibitor exacerbated it. Combined miRNA-21-3p inhibitor with *VEGFA* siRNA weakened the migration effects (Figures 4A and 4B). Tubule and branch formation increased after *CCM2* knockdown. MiRNA-21-3p mimic significantly reduced tubule and branch formation, while miRNA-21-3p inhibitor did not exacerbate it. Combined miRNA-21-3p inhibitor with *VEGFA* siRNA weakened tubule and branch formation effects (Figures 4C and 4D). These results suggested that miRNA-21-3p reduced angiogenesis in *CCM2*-depleted BMEC by regulating VEGFA.

**Figure 4.**
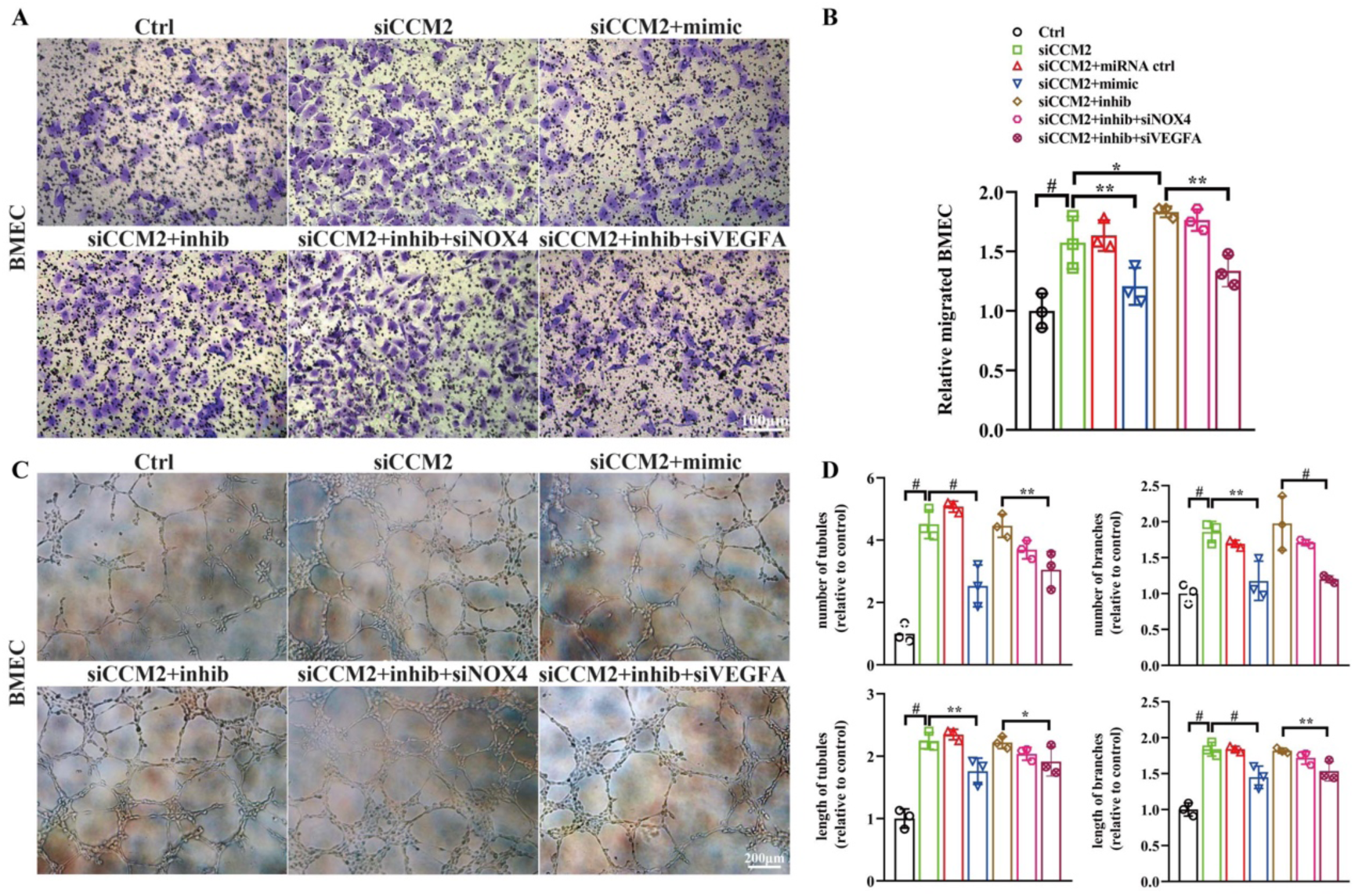
MiR-21-3p inhibits endothelial cell migration and tubule formation by downregulating VEGFA expression in *CCM2*-deficient human brain microvascular endothelial cells (BMECs). A and B, Effect of miR-21-3p mimic and inhibitor on the migration of *CCM2*-deficient human BMECs by Transwell chamber. C and D, Effect of miR-21-3p mimic and inhibitor on the tubule formation of *CCM2*-deficient human BMECs by detecting the number and length of tubules and branches formation. Mean±SD are provided (n=3). *P<0.05; **P<0.01; ^#^P<0.001.

### 3.7. MiRNA-21-3p regulates NOX4 and VEGFA expression in a human pericyte model with *CCM2* gene depletion

MiRNA-21-3p expression in human pericytes decreased after *CCM2* knockdown but could be modulated with miRNA-21-3p mimics or inhibitor (Figure 5A). NOX4 and VEGFA expression levels increased after *CCM2* knockdown. MiRNA-21-3p mimics inhibited this upregulation, while miRNA-21-3p inhibitor intensified it. Combined miRNA-21-3p inhibitor with *NOX4* or *VEGFA* siRNA weakened the upregulation effects (Figures 5A and 5B).

**Figure 5.**
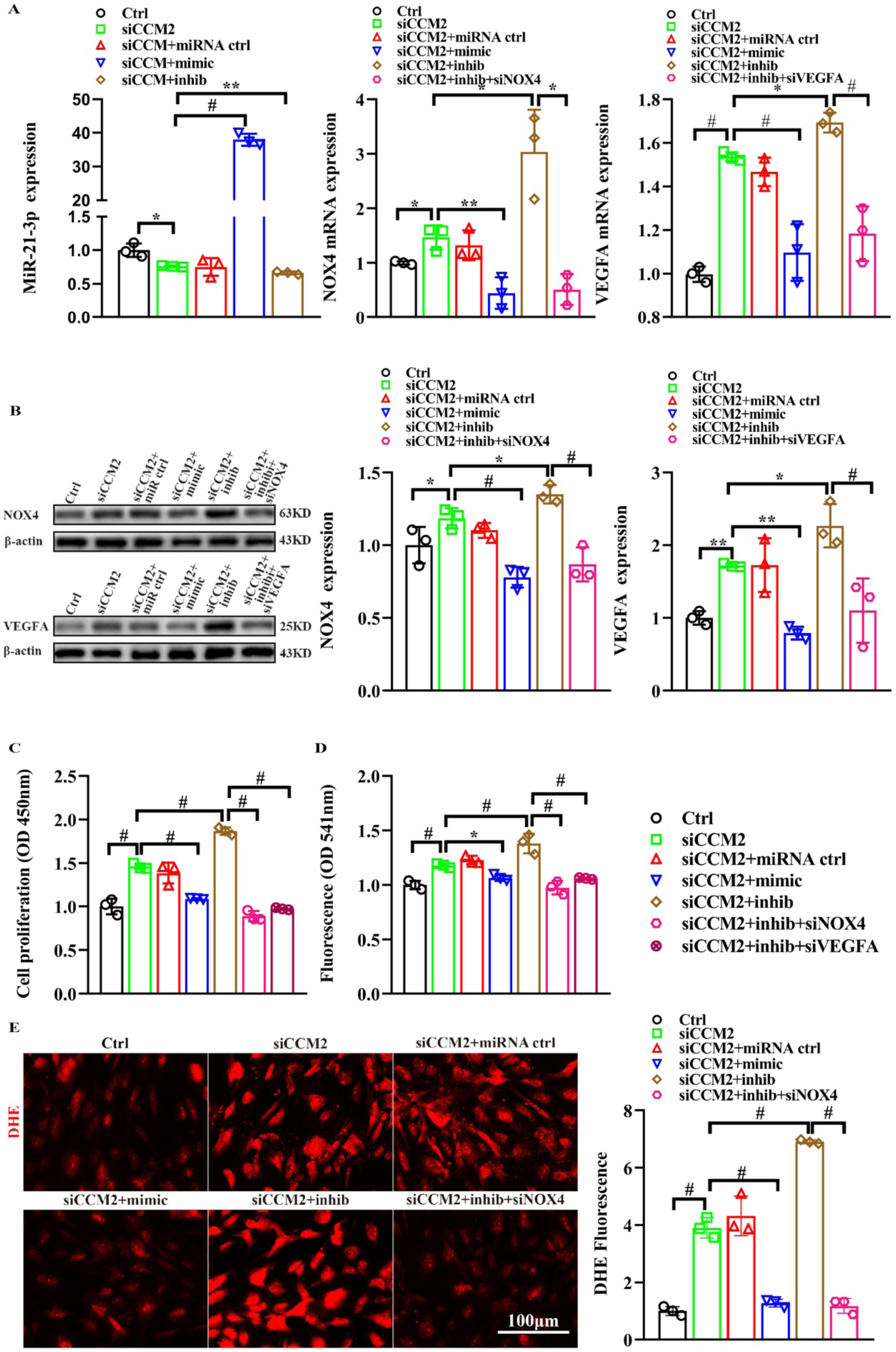
MiR-21-3p inhibits cell proliferation, pericyte permeability, and reactive oxygen species generation by suppressing the expression of the target genes NOX4 and VEGFA in *CCM2*-deficient human pericytes. A, RT-PCR analysis of miR-21-3p, *NOX4* and *VEGFA* mRNA expression in *CCM2*-deficient pericytes transfected with miR-21-3p mimic and inhibitor. B, Western blot analysis of NOX4 and VEGFA in *CCM2*-deficient pericytes transfected with miR-21-3p mimic and inhibitor. C, *CCM2*-deficient pericytes transfected with miR-21-3p mimic and inhibitor were subjected to the CCK-8 assay after transfection. D, Effect of miR-21-3p mimic and inhibitor on the permeability of the *CCM2*-deficient pericytes monolayers to sodium fluorescein by in vitro permeability assay. E, Detection of superoxide by DHE in *CCM2*-deficient pericytes transfected with miR-21-3p mimic and inhibitor. Mean ± SD are provided (n=3). *P<0.05; **P<0.01; ^#^P<0.001.

### 3.8. MiRNA-21-3p inhibits cell proliferation, permeability, and ROS by regulating NOX4 and VEGFA in a human pericyte model with *CCM2* gene depletion

Pericyte proliferation increased after *CCM2* knockdown. MiRNA-21-3p mimic inhibited proliferation, while miRNA-21-3p inhibitor had the opposite effect. Combined miRNA-21-3p inhibitor with *NOX4* or *VEGFA* siRNA weakened the proliferation effects (Figure 5C). Pericyte permeability increased following *CCM2* knockdown. MiRNA-21-3p mimic intervention restrained the rise, while miRNA-21-3p inhibitor intensified it. Combined miRNA-21-3p inhibitor with *NOX4* or *VEGFA* siRNA weakened permeability effects (Figure 5D). These results suggested that miRNA-21-3p attenuated cell proliferation and permeability in *CCM2*-deficient pericytes by regulating NOX4 and VEGFA.

Superoxide anion levels in pericytes increased after *CCM2* knockdown. MiRNA-21-3p mimic inhibited this rise, while miRNA-21-3p inhibitor intensified it. Combined miRNA-21-3p inhibitor with *NOX4* siRNA weakened the upregulation effects. Hydrogen peroxide levels did not significantly increase, suggesting miRNA-21-3p attenuates ROS expression in *CCM2*-deficient pericytes by regulating NOX4 (Figures 5E).

### 3.9. MiRNA-21-3p regulates NOX4 and VEGFA expression in a zebrafish model with *ccm*2 gene knockdown

Different expression levels of miRNA-21-3p, *ccm2*, *nox4*, and *vegfa* were observed in the brain of Tg (Flk1:eGFP; Gata1:DsRed) larvae zebrafish (Supplementary Figure S2). MiRNA-21-3p expression reduced in the zebrafish brain tissue after *ccm2* gene knockdown with MOs, and subsequent increase or suppression was achieved with miRNA-21-3p mimic or miRNA-21-3p MO. The expression levels of *nox4* and *vegfa* in zebrafish brain increased after *ccm*2 gene and miRNA-21-3p knockdown but were inhibited by miRNA-21-3p mimic (Figure 6A).

**Figure 6.**
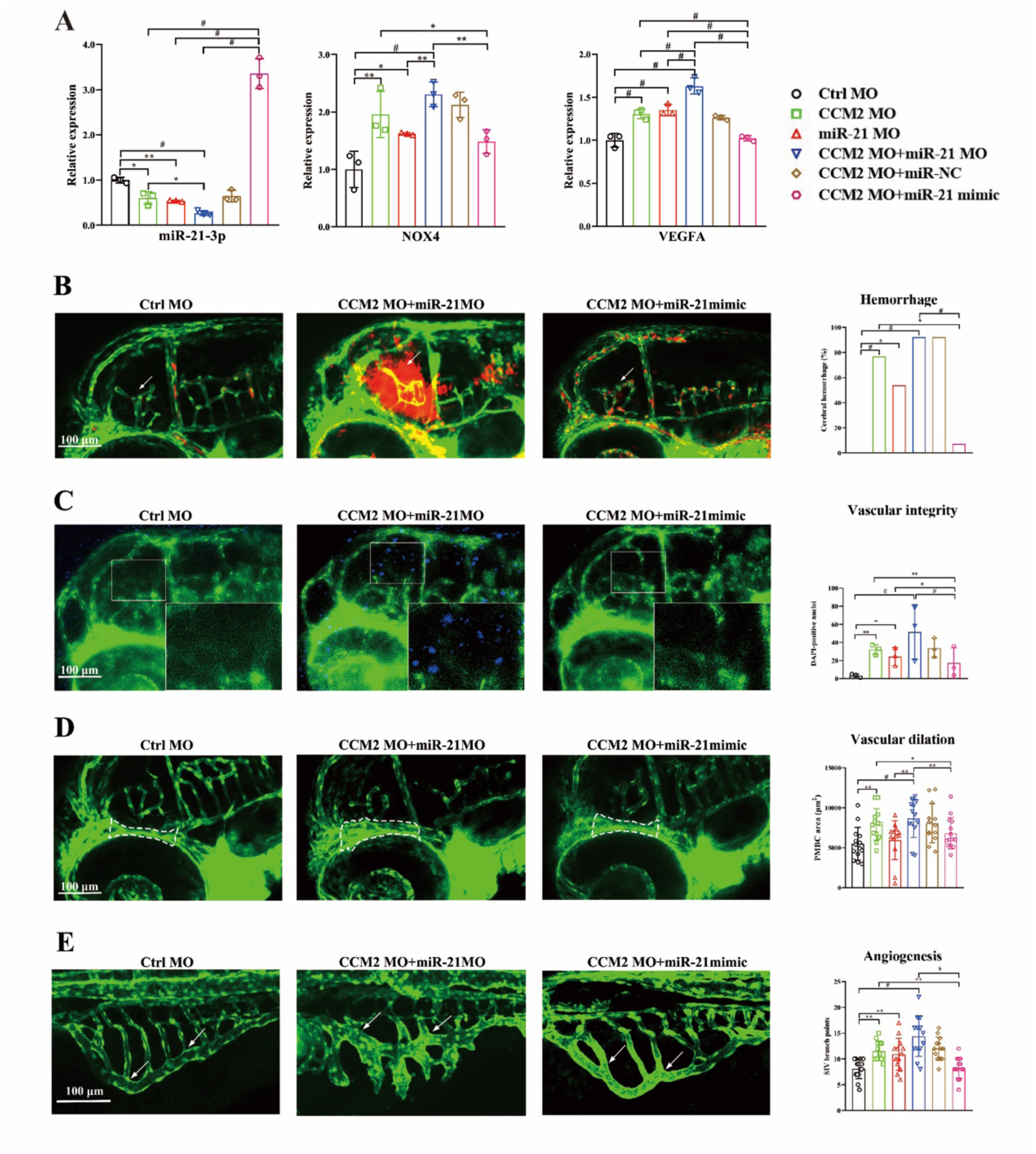
In *ccm2* knockdown zebrafish, miR-21-3p reduces cerebral hemorrhage, vascular permeability, vascular dilation and angiogenesis by inhibiting the expression of nox4 and vegfa. A, RT-PCR analysis of miR-21-3p, nox4, and vegfa expression in *ccm2* knockdown zebrafish with miR-21-3p mimic and miR-21-3p morpholino oligonucleotides (MOs). Three zebrafish embryos were used for each group. B, miR-21-3p mimic decreased erythrocyte leakage and cerebral hemorrhage (arrowheads) in the brain of Tg (Fli1: eGFP; Gata1: DsRed) zebrafish with *ccm2* knockdown. Thirteen to fourteen zebrafish embryos were used for each group. C, miR-21-3p mimic decreased the number of DAPI-positive brain parenchymal nuclei (blue) of Tg (Fli1: eGFP) zebrafish with *ccm2* knockdown. Three zebrafish embryos were used for each group. D, miR-21-3p mimic decreased the areas of primordial midbrain channel (white dotted lines) of Tg (Fli1: eGFP) zebrafish with *ccm2* knockdown. Thirteen to fourteen zebrafish embryos were used for each group. E, miR-21-3p mimic decreased the areas of subintestinal vein (arrowheads) of Tg (Fli1: eGFP) zebrafish with *ccm2* knockdown. Thirteen to fourteen zebrafish embryos were used for each group. Green indicating blood vessels, red indicating erythrocytes and, blue indicating DAPI. *P<0.05; **P<0.01; ^#^P<0.001.

### 3.10. MiRNA-21-3p reduces cerebral hemorrhage, vascular permeability, vascular dilation and angiogenesis in a zebrafish model with *ccm2* gene knockdown

Cerebral hemorrhage phenotype was found after knocking down the *ccm2* and miRNA-21-3p genes. The occurrence of obvious cerebral hemorrhage in our study was classified as the hemorrhage area in the brain of more than 500 µm^2^. Increased miRNA-21-3p expression reduced the zebrafish experiencing bleeding compared to the *ccm2* and miRNA-21-3p knockdown group, as measured by the cerebral hemorrhage rate as well as the hemorrhage area (Figure 6B). These results underscore the crucial role of miRNA-21-3p in mitigating cerebral hemorrhage in zebrafish with *ccm2* gene knockdown.

We found a significant increase of DAPI-positive nuclei of brain parenchymal cells after *ccm2* and miRNA-21-3p gene knockdown. Increased miRNA-21-3p reduced the increased DAPI-positive cell nuclei in zebrafish brains compared to *ccm2* and miRNA-21-3p knockdown group (Figure 6C). These results suggested increased miRNA-21-3p reduced brain vascular permeability in a zebrafish model with *ccm2* gene knockdown.

Our study found a significant increase in the PMBC area after *ccm2* and miRNA-21-3p gene knockdown. Increased miRNA-21-3p suppressed the elevated PMBC area in zebrafish brains compared to *ccm2* and miRNA-21-3p knockdown group (Figure 6D). These results suggested increased miRNA-21-3p attenuated vascular dilation in a zebrafish model with *ccm2* gene knockdown.

Our results also indicated a significantly increased SIV sprouting after *ccm2* and miRNA-21-3p gene knockdown. MiRNA-21-3p mimic significantly inhibited the increase of SIV sprouting compared to *ccm2* and miRNA-21-3p knockdown group (Figure 6E). These results suggested increased miRNA-21-3p inhibited angiogenesis in a zebrafish model with *ccm2* gene knockdown.

## 4. Discussion

In this study, we clarify the association of endothelial and pericyte miRNA-21-3p with cerebral hemorrhage in human CCM lesions, as well as in a human BMEC and pericyte model with *CCM2* gene depletion and a zebrafish model with *CCM2* gene knockdown. MiRNA-21-3p, as a novel therapeutic target, plays a critical role in reducing hemorrhage in CCM disease by regulating its two direct target genes, NOX4 and VEGFA, which are involved in oxidative stress and angiogenesis in both endothelial cells and pericytes. Under the *CCM2* gene-depleted or knockdown state, increased miRNA-21-3p inhibits cell proliferation, permeability, ROS expression, cell migration, and tubule formation by downregulating NOX4 and VEGFA, ultimately leading to a reduction in cerebral hemorrhage, vascular permeability, vascular dilation, and angiogenesis in the zebrafish model. Our study provides further evidence that microRNAs may be potential therapeutic targets for CCM disease.

Our study found that the miRNA-21-3p/NOX4/ROS pathway in *CCM2* gene-depleted BMECs and pericytes plays a crucial role in the cerebral hemorrhage of CCM disease. Increased NOX4 expression promotes excessive ROS production, leading to the dysfunction of BMECs and pericytes, which results in increased vascular permeability and cell proliferation. Upregulated NOX4 was significantly inhibited by intervention with the miRNA-21-3p mimic, resulting in a reduction of excessive ROS generation in the BMECs and pericytes and alleviation of cerebral hemorrhage in a zebrafish model. Previous studies have demonstrated that mutations causing functional loss in the *CCM* gene can lead to various effects, including defects in autophagy, alterations in ROS homeostasis, and increased sensitivity to oxidative stress and inflammation in CCM disease [34,35]. NOX4 is primarily produced by BMECs, with abundant expression in the brain. The pro-inflammatory cytokine TNF-α regulates the expression of NOX4 in BMECs, leading to an increase in ROS and cell apoptosis. The effect of knocking down NOX4 expression or inhibiting NOX4 activity can significantly alleviate oxidative stress and cell apoptosis caused by TNF-α, and has a protective effect on BMECs [36–38]. *CCM1* gene dysfunction in a mouse model and endothelial cells upregulates NOX4 expression, leading to excessive ROS production in endothelial cells, which exacerbates the increased permeability of endothelial cells caused by various inflammatory stimuli and ultimately impairs vascular barrier function [39,40].

Cancer cells exhibit stronger oxidative stress due to elevated levels of NOX and mitochondrial dysfunction compared to normal cells. The level of ROS derived from NOX is significantly increased in cancer cells, while miRNA-21a-3p, miRNA-34a, miRNA-137, and miRNA-99a targeting NOX are significantly downregulated in ROS-driven cancer [41]. MiRNA-21a-3p functions as an endogenous inhibitor of NOX4, which is derived from endothelial cells. Upregulating miRNA-21a-3p can inhibit the activity and expression of NOX4, thereby reducing the excessive generation of ROS induced by NOX4 and contributing to the alleviation of oxidative stress, as well as inhibiting the growth of endothelial cell tumors in mice [23]. Recent clinical studies indicate that tempol may effectively treat symptomatic CCMs by reducing ROS levels [42]. Our findings show that the miRNA-21a-3p/NOX4/ROS pathway regulates functions in BMECs with the *CCM2* gene deletion and similarly affects *CCM2* gene-depleted pericytes, thereby broadening the potential value of microRNA in regulating the oxidative stress mechanism across multiple cell types to reduce fatal cerebral hemorrhage in CCM disease.

Recent studies have shown that emerging therapies, including MAPK/ERK inhibitors, RhoA/ROCK inhibitors, and anti-VEGF therapy, yield effective therapeutic outcomes in preclinical models for arteriovenous malformations (AVMs) and CCMs. Predictive and prognostic biomarkers, such as the candidate biomarker VEGF for AVM and CCMs, could improve the ability to personalize treatment decisions [6,11]. In the *CCM1* gene-deficient endothelial cell, the increased expression of VEGFA results in enhanced cell migration. In a mouse model of *CCM1* gene deficiency, a VEGFA receptor inhibitor reduces the number of CCM lesions, normalizes abnormal vascular permeability, and decreases hemorrhage volume by blocking the VEGF signaling pathway, thereby improving endothelial function and inhibiting angiogenesis [43]. Additionally, in a new CCM model with the *Map3k3* ^I441M^ knock-in mouse, the upregulated VEGFA expression leads to the progression of CCM lesions [44]. These studies imply that microRNA-based targeting of VEGF signaling therapies may have a critical role in the management of cerebral vascular malformations. In cerebral AVM lesions, the expression of miRNA-18a is downregulated in BMECs. MiRNA-18a can attenuate the abnormal proliferation and improve functional normalization of BMECs by reducing the secretion of VEGFA. In a mouse AVM model, administration of miRNA-18a mimics inhibits abnormal angiogenesis, thereby suppressing the progression of AVM lesions [45].

Our findings show that elevated VEGFA expression and decreased miRNA-21-3p expression in the BMECs and pericytes of human CCM lesions are associated with cerebral hemorrhage in CCM disease. The protective effect against hemorrhage with miRNA-21-3p arises from its ability to regulate the target gene of VEGFA, as shown in the *CCM2* gene-depleted BMECs and pericytes model and a zebrafish model with *CCM2* gene knockdown. A previous study has also suggested an interaction between miRNA-21-3p and VEGFA, in which miRNA-21-3p mimics inhibit PI3K/AKT/mTOR signaling by targeting VEGFA expression, thereby suppressing autophagy in bovine ovarian granulosa cells [24]. Recent studies offer new insights into the regulation of the VEGF pathway across multiple cell types in CCM lesions, suggesting it as a potential therapeutic target for CCM. The interaction between endothelial cells and pericytes mediated by VEGFA/VEGFR2 signaling leads to increased angiogenesis and vascular permeability [17]. Furthermore, an increase in the numbers of monocytes, neutrophils, and nature killer cells has been observed in CCM lesions, with enhanced intercellular communication primarily through the VEGF pathway [46]. Our findings that miRNA-21-3p regulates VEGFA in BMECs and pericytes present a potential therapeutic opportunity for reducing cerebral hemorrhage by targeting multiple cells in CCM lesions, with miRNA-21-3p as a target.

Our study has certain limitations. The protective role of miRNA-21-3p in reducing hemorrhage was not assessed in a *Ccm2*^ECKO^, *Ccm1*^ECKO^*, Map3k3* ^I441M^ mouse model of CCM disease. The assessment using these mammalian models is more effective for evaluating the therapeutic effect following the CCM lesion state [9,16,44]. Recent studies suggest that multiple cell types contribute to the pathogenesis of CCM disease [17,46,47]. MiRNA-21-3p may be involved in the cross-talk process between these cells during the development of CCM and its associated bleeding, in addition to the roles of endothelial cells and pericytes observed in our study, which warrants further research to clarify.

## 5. Conclusions

Our study sheds light on the effective role of miRNA-21-3p in inhibiting bleeding in CCM disease through regulating oxidative stress and angiogenesis in BMECs and pericytes. MiRNA-21-3p may be a novel therapeutic target for stabilizing vascular integrity and reducing cerebral hemorrhage in CCM lesions by targeting NOX4 and VEGFA.

## Author contribution statement

Z-SS participated in the design of the present study. All authors participated in the interpretation and collection of the data. X-XG, Z-RH and Z-SS wrote the initial manuscript. Z-SS revised the manuscript. All authors critically reviewed and edited the manuscript and approved the final version.

## Funding statement

This study was funded by the National Natural Science Foundation of China (81873752) and the Science and Technology Planning Project of Guangdong Province (2023B1212060018).

## Conflict of interest statement

The authors declare no conflict of interest.

## Ethics statements

The study was reviewed and approved by the Ethics Committee of Sun Yat-sen Memorial Hospital, Sun Yat-sen University. The animal experiment was approved by the Ethics Committee of Sun Yat-sen University.

## Notes

### Competing Interest Statement

The authors have declared no competing interest.

